# Deep Learning-Based Parameter Estimation for Neurophysiological Models of Neuroimaging Data

**DOI:** 10.1101/2022.05.19.492664

**Authors:** John David Griffiths, Zheng Wang, Syed Hussain Ather, Davide Momi, Scott Rich, Andreea Diaconescu, Anthony Randal McIntosh, Kelly Shen

## Abstract

Connectome-based neural mass modelling is the emerging computational neuroscience paradigm for simulating large-scale network dynamics observed in whole-brain activity measurements such as fMRI, M/EEG, and related techniques. Estimating physiological parameters by fitting these models to empirical data is challenging however, due to large network sizes, often physiologically detailed fast-timescale system equations, and the need for long (e.g. tens of minutes) simulation runs. Here we introduce a novel approach to connectome-based neural mass model parameter estimation by employing optimization tools developed for deep learning. We cast the system of differential equations representing both neural and haemodynamic activity dynamics as a deep neural network, implemented within a widely used machine learning programming environment (PyTorch). This allows us to use robust industry-standard optimization algorithms, automatic differentiation for computation of gradients, and other useful functionality. The approach is demonstrated using a connectome-based network with nodal dynamics specified by the two-state RWW mean-field neural mass model equations, which we use here as a model of fMRI-measured activity and correlation fluctuations. Additional optimization constraints are explored and prove fruitful, including restricting the model to domains of parameter space near a bifurcation point that yield metastable dynamics. Using these techniques, we first show robust recovery of physiological model parameters in synthetic data and then, as a proof-of-principle, apply the framework to modelling of empirical resting-state fMRI data from the Human Connectome Project database. For resting state activity, the system can be understood as a deep net that receives uncorrelated noise on its input layer, which is transformed into network-wide modelled functional connectivity on its output layer. This is consistent with the prevailing conception in theoretical neuroscience of resting-state functional connectivity patterns as an emergent phenomenon that is driven by (effectively) random activity fluctuations, which are then in turn spatiotemporally filtered by anatomical connectivity and local neural dynamics.

## 1 Background

### 1.1 Connectome-based neural mass models in neuroimaging and computational neuroscience

Human brain functioning is characterised by complex spatio-temporal activity patterns, and can be investigated via computational models that describe mathematically how this activity evolves in time. Neuroimaging techniques such as functional magnetic resonance imaging (fMRI) and magneto/electroencephalography (M/EEG), allow noninvasive and holistic measurement of these large-scale activity patterns in human subjects. A dominant theoretical perspective in computational modelling of neuroimaging data is that much of the richness and complexity in the observed activity results from the combination of anatomical network structure and local neural dynamics (Honey et al., 2009; Deco et al., 2013b). Connectome-based neural mass models (CNMMs) have, over the past decade, become one of the principal computational tools used to explore scientific questions in this line of research (e.g. Deco et al., 2013b,a, 2014; Deco and Kringelbach, 2014; Breakspear, 2017; Griffiths et al., 2020). In CNMMs, equations describing the mesoscopic collective neural population behaviour in a large patch of tissue are used to model regional neural activity, such as that measured by average blood oxygenation level-dependent (BOLD) fMRI signal within a brain region or ‘parcel’. CNMM network nodes are interconnected with weights representing the strength of long-range white matter projections (the anatomical connectome), typically parameterized in human brain data using diffusion-weighted MRI (dwMRI) tractography (Hagmann et al., 2008).

To date, CNMMs have been used with moderate success to simulate, and thereby better understand, several characteristics of functional neuroimaging data, including static resting-state fMRI (rsfMRI) functional connectivity (FC; Deco et al., 2013b), dynamic (moving-window) FC (Hansen et al., 2015; Kehoe et al., 2017), state switching (Kringelbach and Deco, 2020; Griffiths et al., 2020), power spectra (Xie et al., 2018), graph-theoretic properties (Vecchio et al., 2017), metastability (Deco et al., 2017), travelling waves (Muller et al., 2016; Roberts et al., 2019), and EEG-BOLD anticorrelations (Schirner et al., 2018; Pang and Robinson, 2018). These studies have employed a combination of mathematical/numerical systems analysis (e.g. bifurcations, attractor landscapes, linear stability) and more direct goodness-of-fit evaluations for comparing empirical and simulated data. The latter process of finding the best set of model parameters, as defined by minimizing some explicitly defined objective function, is variously termed ‘parameter estimation’, ‘parameter optimization’, ‘model inversion’, ‘model fitting’, and ‘data fitting’. These terms shall be treated as largely synonymous in the present paper.

Even though CNMMs have been extensively employed to reproduce the spatially distributed patterns of neuronal activity in numerical simulations, methodologies for CNMM parameter estimation remain relatively underdeveloped. Dynamic causal modelling (DCM; Friston et al., 2003), a related family of techniques that has largely focused on smaller networks and simpler models of neural dynamics than CNMMs, makes effective use of relatively sophisticated Variational Bayesian inference techniques. Unlike DCM however (which has always been principally concerned with parameter estimation), research using CNMMs has focused on characterizations of neural system dynamics at the whole-brain scale, mainly through the use of numerical simulations, and to a lesser extent analytic theory. Where parameter estimation has been employed in CNMM studies, it has usually (with some exceptions, discussed later) been ‘brute-force’ - whereby all possible combinations of some constrained parameter space are evaluated. Whilst the brute-force method is by definition the most robust way of identifying optimal parameters for some defined objective function, it is only viable for extremely low-dimensional parameter spaces, due to the combinatorial explosion introduced by every additional parameter added. The number of parameters that can be feasibly covered with a brute-force approach depends on run time of each parameter combination, the granularity of the parameter space sampling, and the compute resources available for concurrent model runs. CNMMs are often quite computationally expensive to run, as they typically model whole-brain activity, across networks with several hundred nodes, with systems of (often stochastic, often nonlinear, often delay) differential equations, with both sub-millisecond integration time steps and run lengths of tens of minutes (the typical session length of an rsfMRI scan). These factors combine to make a single

CNMM simulation run take quite a long time to execute, even on fairly powerful compute hardware. The number of parameters that can be feasibly covered with the brute-force method for typical CNMMs is therefore usually fewer than five.

Properly deployed, modern optimization techniques are easily capable of vastly superior performance than the brute-force or parameter space exploration approach. In this article, we tackle the problem of CNMM parameter estimation using a number of techniques that are novel or have been rarely used in this context to date. In the next few sections we briefly review at a conceptual level some of the key ingredients of our approach, and then go on to present the details of our methodology and results obtained.

### 1.2 Recasting CNMM parameter estimation as learning with deep recurrent neural nets

#### 1.2.1. Black-Box and Glass-Box Approaches

In this work we approach the problem of parameter estimation for CNMMs, and for physiological models more generally, through the lens of modern machine learning with artificial neural networks (ANNs). Parameter estimation methodologies commonly used to date tend to vary in the level of ‘transparency’ given to the physiological elements of the model being optimized. This can be thought of in terms of more or less ‘black-box’ vs. ‘glass-box’ approaches.

The black-box problem involves, as the name suggests, treating the CNMM as an opaque parameters-in, simulation-out system. For this approach, one would wrap a core CNMM numerical simulator component within one of the great many readily-available black-box parameter optimization frameworks, giving the algorithm no additional information on the class of optimization problem or the mathematical structure inside the box. The advantage of this approach is that it allows the researcher to use the same software implementations, such as those in various widely used computational neuroscience simulator libraries, that they are already familiar with and have expertise in. These implementations are themselves often highly complex and heavily optimized for numerical performance by domain experts, so ‘opening up the box’ may be neither desirable nor feasible.

Although black-box approaches to parameter estimation have the advantage of simplicity, it is widely understood that performance of optimization algorithms can be improved considerably by ‘glassening’ the box in one way or another - giving the algorithm some information about the internal structure, or at least the input-output behaviour, of the system.

Two ways that optimization performance can be enhanced through knowledge of the problem are:

##### 1) Appropriate choice of optimization algorithm

The choice of algorithm may be determined by considerations such as whether the geometry of the objective function or type of problem requires a method that is continuous or discrete, constrained or unconstrained, single or multi-objective, deterministic or stochastic, etc.

##### 2) Jacobians

In the context of parameter optimization, the Jacobian is a matrix of partial derivatives of model parameters with respect to the objective function. Knowledge of the Jacobian dramatically speeds up and improves the convergence of numerical optimization routines. In some cases, the Jacobian can be found analytically, although this is prone to error and relatively rare in computational neuroscience applications, which often use complex sets of equations to model neural activity. More common is to approximate the Jacobian by first linearizing the model equations, and then evaluating the system’s eigenvalues at its fixed points^1^. In other cases the Jacobian can be found numerically, without any specialised mathematical knowledge or reformulation. The traditional way this is done computationally is by numerically approximating the Jacobian using, for example, finite difference techniques. More recently, however, scientific computing tools have begun to make use of automatic (also known as algorithmic) differentiation (AD). AD (or often just ‘autodiff’) is a computational technique that allows implicit application of the chain rule, allowing calculation of derivatives, or ‘gradients’, between variables in a computer program that are separated by long sequences of elementary mathematical operations (see Baydin et al., 2018 for a recent review of AD in machine learning). Critically, in an optimization context, AD allows computation of the derivative of the objective function with respect to the CNMM model parameters, for models with arbitrarily high levels of complexity. AD allows a manifestly glass-box approach to parameter estimation, because it exposes the full set of numerical operations that together define the neural simulation, the observation model, and its associated objective function to the optimization algorithm. Unlike the black-box approach that isolates and separates the CNMM simulator from the optimization machinery, the glass-box approach uses the model equations written out entirely within a dedicated general-purpose numerical parameter optimization (and/or ANN modelling) library such as PyTorch (Paszke et al., 2019), TensorFlow (Abadi et al., 2016), Jax (Bradbury et al., 2018), or STAN (Kucukelbir et al., 2015). We return to the significance of this approach from the point of view of automatic gradient calculation later. First, let us consider the idea of CNMMs and neural simulators more broadly as computation graphs.

#### 1.2.2 CNMMs as Computation Graphs

Computation graphs are software data structures that describe the sequence of numerical operations and input/output relationships along all steps in some computational process. They are specified independently of the actual values of their constituent variables, and play both a schematic and an executional role in computational tasks. A canonical type of computation graph is a deep ANN intended for image classification applications - consisting of states (‘activity’ level at each network node), parameters (e.g. ‘synaptic’ weights between nodes), and equations describing how network activity responds to a given input. Interestingly, from this point of view, CNMMs have a strikingly similar overall architecture to many ANN implementations (recurrent neural networks in particular), consisting as they do of a graph of dynamic ODEs that circulate activation patterns via continuous-valued states, coupled via saturating nonlinearities. Moreover, modern ANN tools not only allow highly flexible selections of off-the-shelf ANN architectures, but also increasingly support the specification of computation graphs directly from arbitrary sets of differential equations. Indeed, it has become clear in recent years that combining conventional ANN architectures and algorithms with differential equation solvers and frameworks opens powerful new directions not only for machine learning itself (e.g. with ‘neural ODEs’; Chen et al., 2018), but also for theoretically-oriented scientific fields such as physics (Raissi et al., 2019). In neuroscience, the above developments have enabled progress both in the ML→Neuro direction - identifying structural patterns in trained deep nets and searching for comparable patterns in the human brain (Güçlü and van Gerven, 2015), and also in the Neuro→ML direction - identifying structural patterns in human brain networks and asking whether they facilitate performance of neural nets on cognitive tasks (Suárez et al., 2021).

### 1.3 Present work

In summary, there are a variety of interesting and practically useful intersections between modern machine learning computational architectures and optimization frameworks, the emerging paradigm of neurophysiological modelling with CNMMs, and physiological modelling more generally. Motivated by this, we have developed a novel model optimization framework that improves on previous work in this area in a number of ways. We demonstrate this approach for a familiar class of CNMMs for resting-state fMRI activity, using a widely-used neural mass model (the two-state RWW equations; Deco et al., 2013b, 2014) and associated haemodynamic model (Friston et al., 2000) to describe regional neural dynamics at every brain region or network node in a whole-brain connectivity graph. Our focus is principally on rsfMRI FC patterns, although the methodology is quite generic, and in other recent work (Momi et al., in prep) we have also found it performs equally well. The framework’s effectiveness is largely owed to a combination of an efficient time-windowed batching and epoching scheme, autodiff-computed gradients, and some selected parameter constraints.

In the following sections we first detail the modelling approach, including equations defining the neural dynamics, objective function, and numerical bifurcation analyses that are used to support parameter constraints. We then present two sets of modelling results, demonstrating i) accurate recovery of parameters in a synthetic dataset with known ground truth, and ii) model-based single-subject analyses of human connectome project (HCP) rsfMRI data. We then conclude with a discussion around the innovations, implications, and limitations of the approach.

## 2 Methods

The chief contribution of this article is the introduction of a novel computational framework for CNMM parameter estimation. A schematic of the general approach is given in Figure 1. The following sections outline the associated neuroimaging data analysis details, followed by the mathematical theory and implementation details for both the neurophysiological model and the optimization algorithm.

**Fig. 1.**
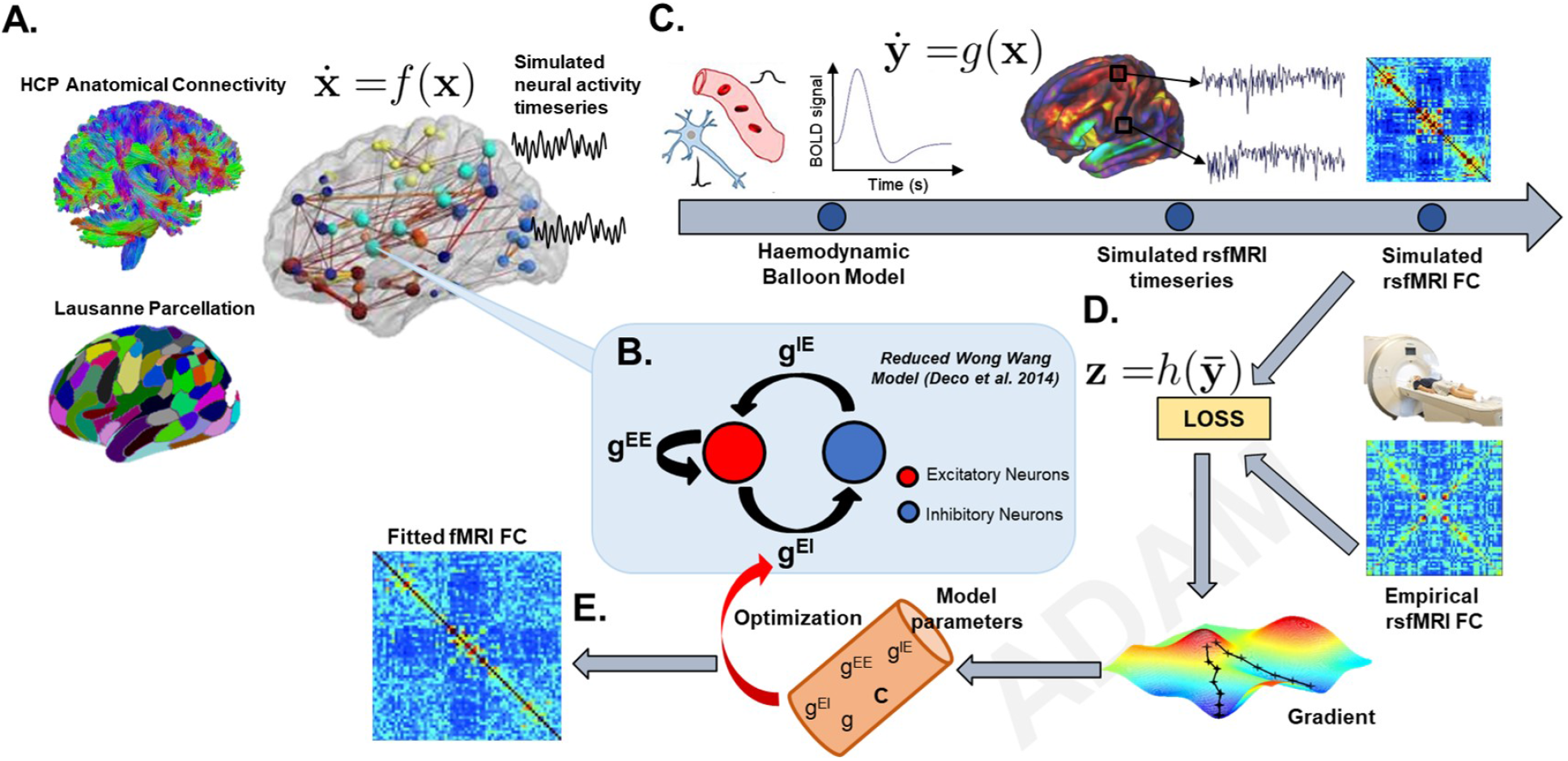
Schematic of CNMM modelling approach. We model rsfMRI data by specifying a deep recurrent neural network having RWW network node dynamics, anatomical connectome network structure, and a haemodynamic observation model, within an autodiff-capable Python programming environment (PyTorch). **A)** Network nodes are defined from the grey matter parcellation, and network edge weights between regions are defined from normalized, re-scaled DWI tractography streamline counts. **B)** Each network node comprises an excitatory and inhibitory neural population, coupled by local gain parameters *g*^*EE*^,*g*^*EI*^, *g*^*IE*^, connected between nodes by the connectome weights *C* and global scaling *g*. **C)** A haemodynamic balloon model converts the neural population activity to simulated regional BOLD fMRI time series, and a whole-brain FC matrix is computed from the Pearson correlation between parcel time series. **D)** Complexity-penalized comparison of simulated to empirical rsfMRI FC matrices informs the objective function (‘LOSS’), which is iteratively fed back into the network, and the AD-computed gradient between the objective function and model parameters is used by the ADAM optimization algorithm to update the model parameters. The model is run with new parameters for a short period (30s simulated time), and the process is repeated until **E)** convergence, giving the final simulated FC matrix and estimated model parameters.

### 2.1 Implementation

There is a large and rapidly-growing array of computational tools used for modelling and optimization in both the academic and commercial sectors. These include the PyTorch (Paszke et al., 2019), TensorFlow (Abadi et al., 2016), Jax (Bradbury et al., 2018), and STAN (Kucukelbir et al., 2015) libraries, amongst others. Here we use PyTorch for the neural network modelling and model-fitting work presented in this paper, which is comparable in most relevant features to the other libraries listed above, with the notable advantage over some of a large, dynamic, and rapidly growing user and developer community. Full code for reproduction of the data analyses and model fitting described in this paper is freely available online at github.com/griffithslab/dl-paramest-for-neurophys-models. For clarity, this repository also includes documentation detailing where and how each of the 40 equations given below appear in the code. All models and analyses were run with Python 3.7 on quad-core processors running Ubuntu 18.0. Full Python library version specifications are given in the repo’s dependencies file.

### 2.2 Neuroimaging data analyses

The CNMM methodology requires two principal data types: a) measured physiological activity (typically in time series or FC matrix form), and b) an anatomical connectome. The aim is to simulate the former using the network structure from the latter, plus an appropriately chosen model of neural dynamics. Here our focus is on modelling resting-state fMRI data, although the general approach is also easily applied to EEG, MEG, and other data types. We have explored the application of our modelling approach on several neuroimaging datasets, but in the interests of space, focus, and simplicity, we focus in this paper on one: the Washington University-Minnesota (WU-Minn) Human Connectome Project (HCP) consortium data (Van Essen et al., 2013; Glasser et al., 2013).

Neuroimaging data analyses were conducted in Python using Nilearn (Abraham et al., 2014), Dipy (Garyfallidis et al., 2014), Nipype (Gorgolewski et al., 2011), and other tools in the Neuroimaging-In-Python ecosystem (github.com/nipy). Anatomical connectivity matrices for 50 randomly chosen subjects were computed from HCP dwMRI data using deterministic tractography. Full details on the local tissue model and dwMRI and tractography analysis pipeline are given in a previous article (Griffiths et al., 2020), and the original raw data (Van Essen et al., 2013) is publicly accessible at https://db.humanconnectome.org. The parcellation used for both dwMRI and fMRI analyses was the scale-1 (83 node) ‘Lausanne 2008’ parcellation (Hagmann et al., 2008; Daducci et al., 2012), which consists of 68 cortical and 15 subcortical regions. These parcels were defined initially on the Freesurfer cortical surface, and converted to nifti volumes and resliced to the native-space EPI images for fMRI data analysis. Using this parcellation, regional time series were extracted from preprocessed rsfMRI nifti volumes using tools from the Nilearn library, with noise covariates from CSF signal, white matter signal, and motion parameters regressed out. FC matrices were calculated as the Pearson correlation between regionally averaged time series from all pairs of parcels.

In addition to these full neuroimaging data analyses, we also used the model to generate synthetic rsfMRI time series, for which the ‘ground truth’ model parameters are known. In these synthetic data analyses, we employed a different (but related) anatomical connectivity matrix: the 68-region ‘Hagmann connectivity’ matrix, first described in the seminal studies of Hagmann et al. (2008) and Daducci et al. (2012), and used extensively in modelling work since then (e.g. Deco et al., 2013b, 2014; Robinson, 2012; Mehta-Pandejee et al., 2017). As in these latter studies, for this set of synthetic data simulations and analyses (described in Results section 3.1) we did not ourselves run dwMRI tractography to generate the connectivity matrices, but instead used the matrices provided by the above authors (Deco et al., 2013b; Hagmann et al., 2008).

### 2.3 Ethics Statement

The computational modelling work undertaken in this study was approved by the Centre for Addiction and Mental Health (CAMH) Research Ethics Board (REB number 022/2020). The modelling The human neuroimaging datasets used in this study were collected by other research teams as part of the Human Connectome Project (Van Essen et al., 2013), with all participants providing written informed consent prior to data collection, and was approved by the Washington University Institutional Review Board (IRB number 201204036).

### 2.4 Neurophysiological model

We modelled the dynamics of whole-brain neural activity with the aim of reproducing the main features of the individual subjects’ FC matrices. As shown in Figure 1, a neural mass model describing the temporal dynamics of regional brain activity was placed at each node of the anatomical parcellation, and coupled to other brain regions according to the weights given by the anatomical connectivity matrix. The regional neural dynamics were then fed into a haemodynamic model describing the conversion of neural activity to BOLD signal fluctuations. Pearson correlation FC matrices were then computed from this simulated regional BOLD activity, exactly the same as with the preprocessed regionally averaged empirical BOLD signals. A general mathematical description of this is as follows:

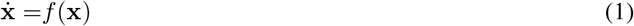

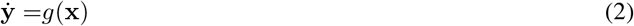

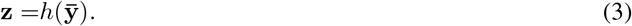

Here, the local neural dynamics (*f*) determine the time-evolution of the state vector **x** = [*x*_1_ … *x*_*N*_] of regional neural activity across *N* network nodes. The function *g* is a so-called haemodynamic *forward* (also known as *measurement* or *observation*) model that maps neural activity on to the BOLD signal time series, as measured by fMRI. Finally, *h* applies the Pearson correlation coefficient, acting over a temporal window 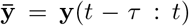 of the time series **y**,where *τ* = 30*s*. In the following we give the details of the functions *f* and *g* for neural activity and haemodynamics, respectively. Subsequent sections give details on construction of the objective function used in parameter estimation using *h* and **z**.

#### Neural dynamics

To describe mesoscopic population activity at each region in the whole-brain network, we use the equations derived in Deco et al. (2013b, 2014), which are variously termed in the literature the ‘Dynamic Mean Field’, ‘Wong-Wang-Deco’ (WWD), and ‘Reduced Wong-Wang’ (RWW) model. Here we use the latter appellation. Specifically, we use the *two-state* or *E-I* version of the RWW equations. In this model, each brain network node consists of two neural masses: one describing the average behavior of an excitatory neural subpopulation, and the other the behaviour of an inhibitory neural subpopulation. Activity in each of these subpopulations at a given brain network node *i* ϵ [1…*N*] is described by the unitless state variables *E*_*i*_ / *I*_*i*_ representing synaptic activity or ‘gating’ levels, as well as auxiliary variables 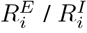 (population firing rates) and 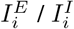 (synaptic currents). These evolve according to the following coupled nonlinear stochastic differential equations:

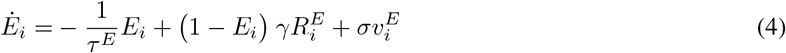

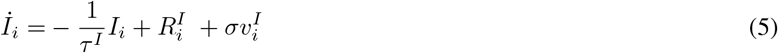

where *τ*^*E*^ and *τ*^*I*^ are the decay times of the excitatory and inhibitory synapses, respectively, *γ* is a kinetic parameter, and 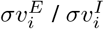 are uncorrelated Gaussian noise processes with mean 0 and standard deviation *σ*. The principal input to *E*_*i*_ and *I*_*i*_ are the population firing rates 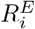 and 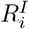, which are expressed as functions of the input currents 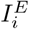 and 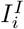 as follows:

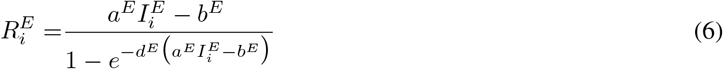

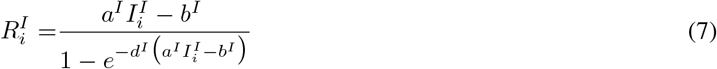

Parameters *a*^*E*^, *a*^*I*^, *b*^*E*^, *b*^*I*^, *d*^*E*^ and *d*^*I*^ govern the specifics of the conversion of input currents to population firing rates (see Table S1 for further info). The two input currents are computed as follows:

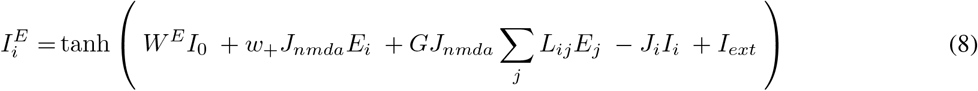

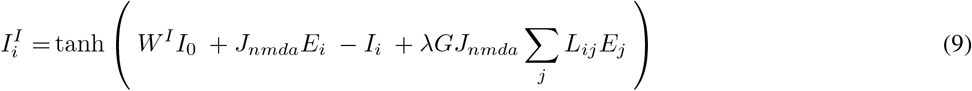

where *I*_*ext*_ encodes external stimulation, and is set to 0 when simulating resting state activity. *I*_0_ represents a constant external input, scaled by parameters *W*^*E*^ and *W*^*I*^ for the excitatory and inhibitory populations, respectively. *L*_*ij*_ denotes the elements of the connectivity Laplacian, defined as *L* = *D − C*, where *C* is the (log-transformed and unit-normalized) tractography-derived connectivity (adjacency) matrix that gives the connection strength between network nodes *i* and *j*, and *D* is the diagonal matrix of node degree (i.e. the row sums of *C*). The term Σ_*j*_ *L*_*ij*_ *E*_*j*_ thus encodes the total summed input to node *i* from all other *j* nodes in the network. The parameter *λ* allows the removal of long-range feedforward inhibition (Deco et al., 2014) when set to 0, which is what was done here (although we retain the term in the equations here for completeness and potential exploration in future work). Parameters *J*_*nmda*_ and *J*_*i*_ represent the value of the excitatory synaptic coupling and the local feedback inhibitory synaptic coupling, respectively, while parameters *w*_+_ and *G* scale the local and long-range excitatory couplings, respectively.

An interesting and important feature of the RWW model is that it is one of the few widely used neural mass models that is directly derived, via mean field reduction techniques, from a ‘lower level’ mathematical description of individual (conductance-based leaky integrate-and-fire) neurons. Building on original work of Wong and Wang (2006), this derivation was first done in the CNMM context for the one-state (slow excitatory population only) model by Deco et al. (2013b), and subsequently for the two-state (excitatory and inhibitory population) model in Deco et al. (2014), which give the equations used here. This two-state RWW model is similar in many respects to the classic equations of Wilson and Cowan (1972), who famously introduced the notion of predator/prey-like excitatory/inhibitory population interactions as a generic description of cortical tissue. In both Deco et al. (2013b) and Deco et al. (2014), the authors also derive further reductions, arriving finally at a linearized algebraic expression for neural population covariance structure - a result that is of considerable importance for the broader theory of CNMMs, although not one we shall be considering further in the present paper.

In addition to deriving and introducing the two-state RWW model, Deco et al. (2014) also introduced an iterative algorithm designed to keep the synaptic current terms within a specific biologically-motivated range. Here we employ a different but related constraint: we squash the input current variables in Equations 8 and 9 using a tanh function. This is a more mathematically well-behaved way of specifying variable limits than explicit constraints or iterative algorithms (it is differentiable, for example), and we have found improves model fit performance considerably.

The notation above follows closely that of Deco et al. (2014). In this paper we also adopt the following alternative notation for several commonly discussed terms:

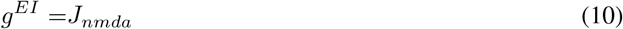

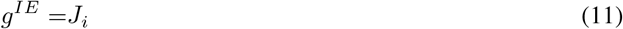

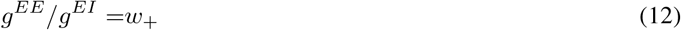

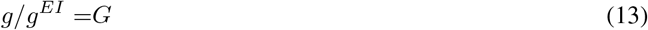

which define the within-node excitatory-to-inhibitory, inhibitory-to-excitatory, and excitatory-to-excitatory synaptic gains (*g*^*EI*^, *g*^*IE*^, *g*^*EE*^, respectively), as well as the long-range global coupling *g*. This new notation is convenient because it is more uniform and follows general conventions in relation to connectivity weight subscripts. We will eventually fit these four parameters by comparing the results of simulations of the model to empirical data.

#### Haemodynamics

The forward model function *g*(**x**) models the transformation of the state variables of the neural mass model into a quantity that can be compared to the experimental BOLD signal obtained by fMRI. Following Deco et al. (2014), for this we use here the Balloon-Windkessel model (Friston et al., 2000), which captures much of the known biophysics of BOLD signal generation.

In the Balloon-Windkessel model, the vascular bed in a small volume of tissue is modelled as an expandable venous compartment. The output of the capillary bed (volume flow rate into the tissue *F*_*i*_(*t*), in units of ml/s) is specified to be a function of time, and drives the system. Blood volume changes happen primarily in the venous compartment. The generated BOLD signal is a nonlinear function of normalized venous volume *V*, normalized total deoxyhemoglobin content *Q*, and resting net oxygen extraction fraction by the capillary bed *O*_*i*_, where rates of change of each variable as the Balloon model expands and contracts are given by

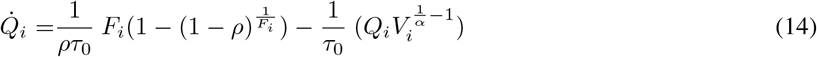

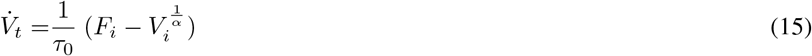

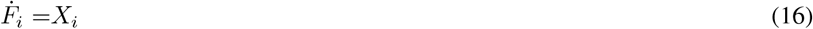

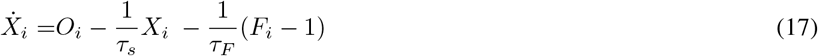

with rate constants *ρ*, transit time *τ*_0_, stiffness parameter *α*, signal decay *τ*_*s*_, and autoregulation *τ*_*F*_ (see Table S1 for nominal values). 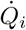 is defined using flow rate multiplied by the concentration, representing the difference between the delivery of deoxyhemoglobin into the venous compartment and the deoxyhemoglobin expelled with the extraction function *O*_*i*_ (defined as 1 – (1 − *ρ*)^1*/F*^) across transit time *τ*_0_. The rate of change of the volume 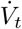 is defined as the difference between inflow *F*_*in*_ and outflow *F*_*out*_ (or 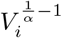) divided by the transit time *τ*_0_. Differentiating 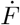 one more time, we obtain 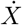 which can model the flow between different steps of the model. The BOLD signal is then given by

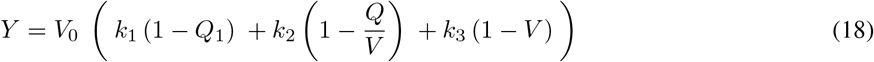

where *V*_0_, *k*_*i*_, *i* = 1, 2, 3 are constant coefficients.

#### Synthetic data generation

For the first round of simulations and analyses reported in this paper, we generated synthetic data using the above equations and known values for model parameters, and tested the ability of the optimization algorithm (see below) to accurately recover the correct parameter values. Importantly, numerical integration of the neural and haemodynamic model equations was run with the same PyTorch code implementation used for model fitting. A set of ‘synthetic subject’ anatomical connectivity (*C*) matrices were generated by adding a small amount of random noise to a template, for which we used the 66×66 tractography-derived matrix from Hagmann et al. (2008) and Deco et al. (2013b). Unlike the *C* parameters, which thus varied slightly across the synthetic subject group, the target neural model parameters *g, g*^*EE*^, *g*^*IE*^, and *g*^*EI*^ were set uniformly to the nominal values listed in Table S1. For each synthetic subject we generated whole-brain regional time series simulations, and these were in turn used to generate a distribution of synthetic simulated rsfMRI FC matrices. Parameter estimation was then run on these synthetic rsfMRI time series, identically as with the empirical data, with randomly selected state variable initialization values. Results of these analyses are shown in terms of distributions over the 50 synthetic subject model runs in their parameter estimates and their parameter estimate errors.

### 2.5 Bifurcation analysis

Bifurcation analyses allow us to define and study qualitative changes in the dynamics of a system that emerge as a function of one or more parameter values, and can be used to summarize some aspect of the dynamical regime at that point in parameter space. These qualitative changes can be extremely useful for understanding the system’s overall behavioural repertoire, and under what conditions a given type of dynamics is expected, guaranteed, or prohibited.

We used bifurcation analyses of the RWW equations to characterize better its dynamical properties, and so doing identify further useful constraints on parameter values during model fitting. The dynamics of a single RWW node without network couplings (c.f. Eqns 4-9) is given by

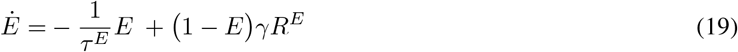

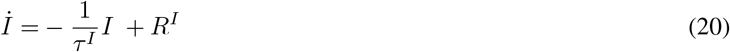

where *R*^*E*^ and *R*^*I*^ are calculated the same as in Eqs. 6-7, and the two input currents are

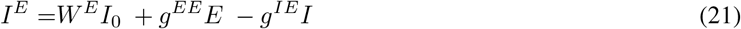

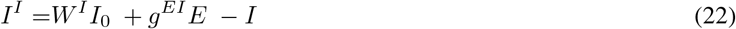

We are interested in dynamic regime changes across the parameters: *I*_0_, *g*^*EE*^, *g*^*IE*^ and *g*^*EI*^. All of these parameters are within the range [0, 2]. For each set of parameters above, using the `fsolve` function from the Python package `scipy.optimize` with multiple random initial parameters, we calculated all the solutions of

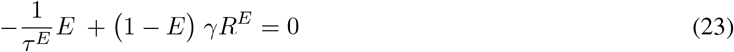

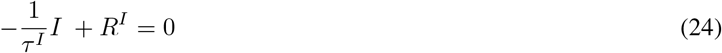

At each solution, the Jacobian was calculated to obtain the corresponding eigenvalues. The number of solutions and the eigenvalues associated with each solution were evaluated, in order determine which dynamic regime those parameters lie within. Within the search scope used, only two regimes were found: a monostable (single fixed point) and a bistable (two fixed points) regime. A subcritical Hopf bifurcation occurs as the monostable region gives rise to the bistable region, where a stable state of steady growth coexists with an oscillatory regime. Here we are interested in the critical parameter values that place the system at the edge of this bifurcation, as this location of parameter space has been identified previously to achieve the best fits with empirical data and most neurobiologically realistic dynamics (Deco et al., 2013b, 2014). To do this we used an iterative approach. First, the critical value of *I*_0_ is calculated through a 1-dimensional parameter search. Then, using that value as a fixed value for *I*_0_, the critical value of *g*^*EE*^ is calculated in a similar manner as for *I*_0_, using a similar search across parameter space. Lastly, both *G*_*io*_ and *g*^*EI*^ are varied, fixing the *I*_0_ and *g*^*EE*^ to the above-determined values. The coloured areas in Figure 2B show the dynamic regimes on the plane spanned by *g*^*IE*^ and *g*^*EI*^. The area labelled *I* represents the fixed point regime, and the area labelled *II* the bistable regime. As is clear from this figure, the boundary between these two regimes reliably takes the form of a decaying exponential. Noting this, we introduced into the optimization algorithm (c.f. Eq. 41) a polynomial function as a joint prior on *g*^*IE*^ and *g*^*EI*^, which serves to ensure that the values of these two parameters are jointly sampled from the critical boundary.

**Fig. 2.**
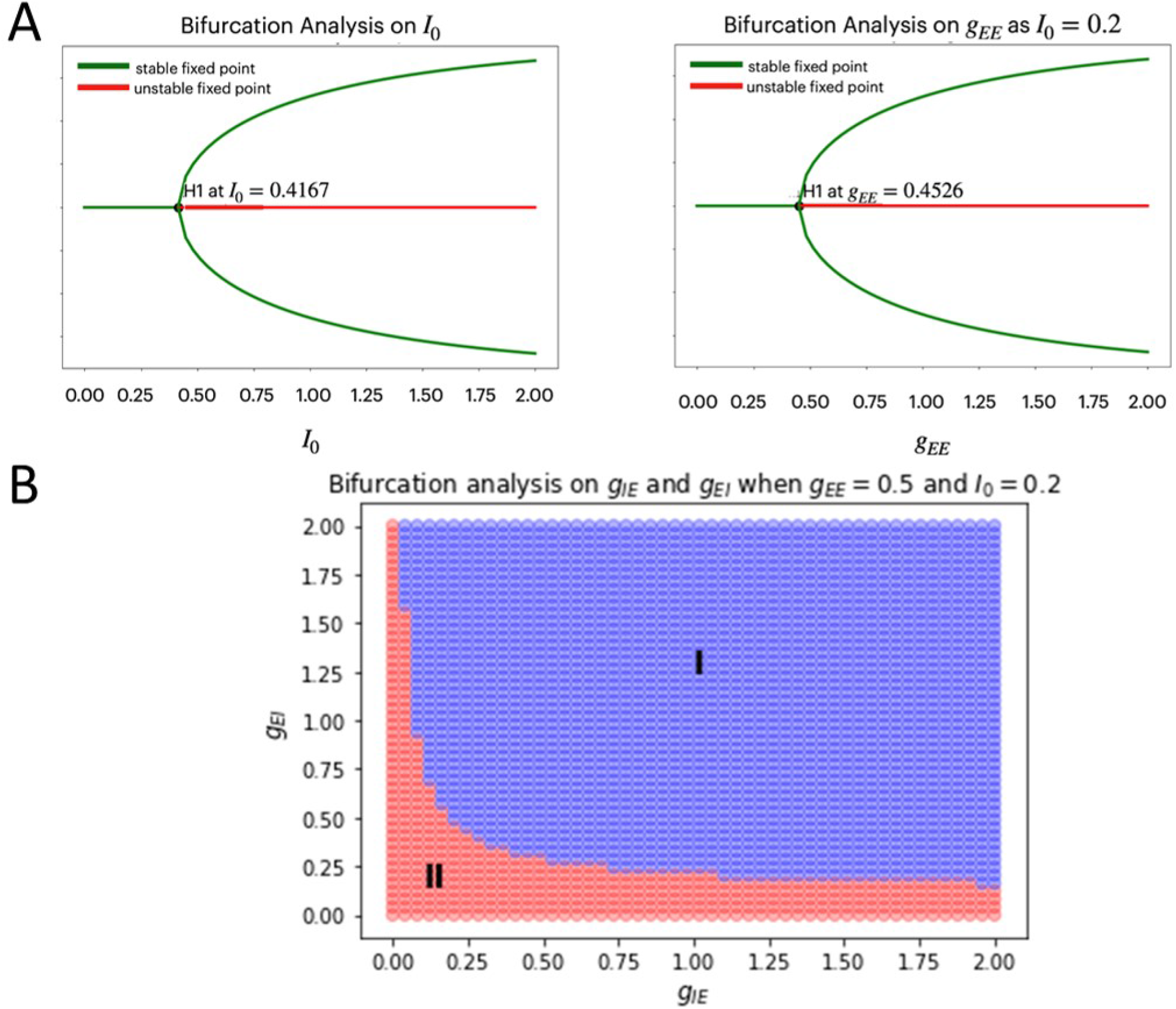
Bifurcation analysis. We used bifurcation analyses of the 2D system describing an isolated RWW node to identify useful parameter constraints. **A)** Bifurcation diagrams showing the bistability associated with the *I*0 and *g*^*EE*^ parameters. When both parameters are varied, two stable points emerge (black dots, labelled H1) at the intersection of the stable and unstable fixed points across potential values for the fixed current *I*0 and the excitatory-excitatory coupling strength *g*^*EE*^ parameters. **B)** After identifying the critical points for *I*0 and *g*^*EE*^ independently, we fix those parameters near to their critical points, and construct a joint bifurcation diagram for inhibitory-excitatory *g*^*EI*^ and excitatory-inhibitory *g*^*IE*^ coupling strengths. This analysis identified two regimes - a fixed-point (I) and an oscillatory (II) regime. Since the separatrix between these two domains of parameter space takes the form of a decaying exponential, we can define a 2D polynomial spline function of *g*^*IE*^ and *g*^*EI*^ that sits close to the border between zone I and zone II. Later, this is polynomial spline is used to define a joint prior density over these two parameters.

### 2.6 Parameter optimization

The brain network model (neural equations 4-13 and haemodynamic equations 14-18) can be understood as a form of recurrent ANN, that uses the parameters *C*_*ij*_ (which in turn define *L*_*ij*_), *g, g*^*EE*^, *g*^*IE*^ and *g*^*EI*^ as its weights. Of these, the *C* parameters in particular are directly analogous to the free parameters of an ANN, determining individual connection strengths between network nodes. The *g* parameters, in contrast, have a more global effect, since they scale the connection weights between all nodes in the network concurrently. All other physiological and haemodynamic model parameters are treated as fixed, and therefore not relevant here. We use a machine learning-based optimization approach to estimate those model parameters by fitting the model-generated activity to empirical functional neuroimaging data recordings.

#### Optimization of model parameters using ADAM

In order to fit the brain network model to the empirical neuroimaging data dynamically, we divide the rsfMRI BOLD time-series into 30s non-overlapping windows (*batches*), as shown in Figure S1. In each batch, the simulated BOLD time series are generated using our model with the currently estimated model parameter values, and the cost is calculated to guide the model parameter updates for the next batch. Our aim is to find optimal parameters *g*^*EI*^, *g*^*IE*^, *g*^*EE*^, *g*, and *C*_*ij*_ in Equations 6-10, to minimize the objective function (defined below at each batch). We used the ADAM algorithm (Kingma and Ba, 2014), which is natively supported in PyTorch, to control the parameter update schedule during model optimization. ADAM is an adaptive learning rate optimization algorithm designed specifically for training deep neural networks, that can be understood as a combination of RMSprop (Tieleman and Hinton, 2012) and Stochastic Gradient Descent (SGD) with momentum. It uses the squared gradients to scale the learning rate (like RMSprop), and it takes advantage of momentum by using the moving average of the gradient instead of gradient itself (like SGD with momentum). Thus, ADAM is an adaptive method that computes individual learning rates by estimating the first and second moments of the gradient of the model parameters with respect to the objective function, and using this to adapt the overall learning rate. The following is a brief summary of the ADAM algorithm:

The *n*-th moment of a random variable *X* is defined as the expected value of that variable to the power of *n*:

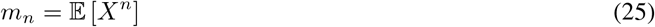

The first moment is the mean, and the second moment is the uncentred variance (i.e. the mean is not subtracted during calculation of the variance). To estimate these moments, ADAM utilizes exponentially moving averages:

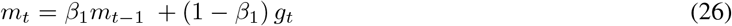

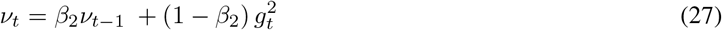

where the estimates of the first and second moments *m*_*t*_ and *v*_*t*_ are moving averages at the *t*-th batch, *g*_*t*_ is the gradient on the current batch, and hyperparameters *β*_1_ and *β*_2_ are set to default values of 0.9 and 0.999, respectively. The vectors of moving averages are initialized with zeros at the first iteration. The estimators are biased towards zero.

Now we need to correct the estimator, so that the expected value is the intended quantity. This step is usually referred to as bias correction. The final formulas for our estimator will be as follows:

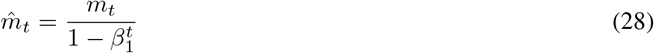

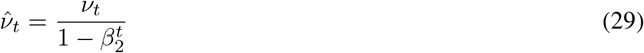

These are the bias-corrected estimators for the first and second moments corrected with bias. The moving averages are used to scale the learning rate individually for each parameter. The way this is done in the ADAM algorithm is very simple: to perform a weight parameter update we take

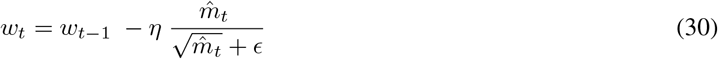

for the model parameters *w* and step size *η*.

#### Objective function

Following other authors, we choose an objective function for the ADAM algorithm based on Variational Bayes that implements a complexity-penalized goodness-of-fit, using the free energy approximation to the log model evidence (Beal, 2003; Friston et al., 2003; Hashemi et al., 2020). Let *X*^*k*^ represent the state (i.e. RWW CNMM state variable vector) at batch *k, y*^*k*^ represent BOLD signals at the batch *k*, and Θ denote all model parameters. We consider the general density

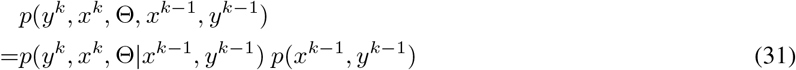

where

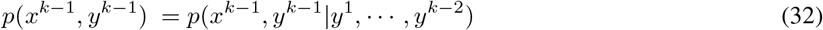

By integrating the state *X* out of Equation 31 we obtain the following compact form for the generative model at batch *k*:

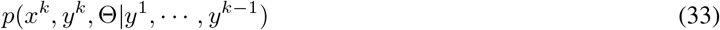

from which we obtain

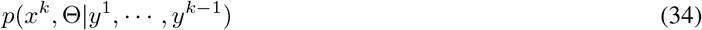

This is the quantity of interest to us, because it describes the posterior probability of the time-dependent states *x*^*k*^ and time-independent parameters Θ.

Variational Bayesian (VB) inversion determines the posterior distributions *p* (*x*^*k*^, Θ|*y*^1^, …, *y*^*k*−1^) by maximizing the log model evidence (LME). The LME corresponds to the negative surprise about the data, given a model, and is approximated by a lower bound, the negative free energy. Detailed treatments of the general principles of the VB procedure can be found in numerous papers (Beal, 2003; Friston and Stephan, 2007). The approximations inherent in VB enable a computationally efficient inversion scheme with closed-form single-step probability, that updates from trial to trial. In particular, VB can incorporate the so-called mean-field approximation, which turns the joint posterior distribution into the product of approximate marginal posterior distributions. Following on from Equation 34,

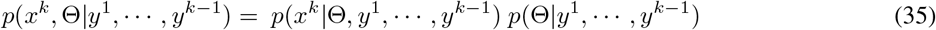

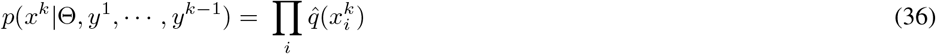

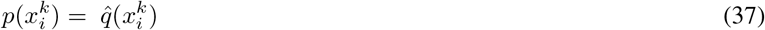

Based on this assumption, the variational maximization of the negative free energy is implemented in a series of variational updates for each state of the model separately. Equation 36 represents the mean-field assumption (according to which the posterior is factorized), while Equation 37 reflects the fact that we assume a fixed form *q* (·) for the approximate marginal 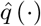. We make minimal assumptions about the form of the approximate posteriors by following the maximum entropy principle: given knowledge of, or assumptions about, constraints on a distribution, the least arbitrary choice of distribution is the one that maximizes entropy (Jaynes, 1957).

Thus, we can define the objective function as:

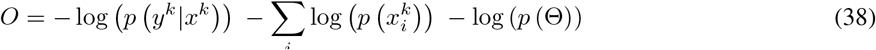

We define

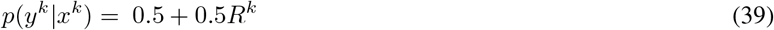

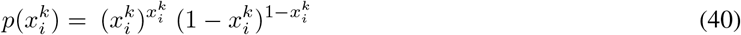

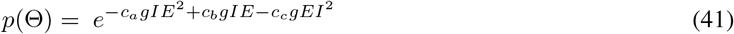

where *c*_*a*_, *c*_*b*_, *c*_*c*_ are hyper-parameters to be fitted, and *R* is the Pearson correlation between simulated and empirical BOLD in the *k*th batch.

Note that Equation 40 is maximizing the entropy of the simulated states on *E* and *I*, which pushes the system towards the bistable regime. Equation 41 is making sure that model parameters are changing around the boundary of the two system regimes (fixed point and bistable), as indicated in the bifurcation analysis of Section 2.4.

## 3 Results

In the following we report the results from application of our model to the cases of i) recovery of known parameters by fitting synthetically generated data, and ii) estimation of parameters from empirical HCP rsfMRI data.

### 3.1 Fit results to synthetic data

We first tested the model fitting scheme’s ability to recover known model parameters, by fitting synthetic data generated by integrating the neural and haemodynamic model equations, as described in Section 2.3. Results from these analyses are shown in Figure 3. At the top of this figure are shown the model-estimated and synthetic ‘empirical’ anatomical connectivity (*C*_*ij*_) and FC values as scatter plots (panel A) and as matrices (panel B) for a single exemplary synthetic subject. The objective function in Equations 38-41 operates on the FC matrix, and so the SC parameters should be understood as being learned though the structure present in the FC. The model fit here is excellent, with a Pearson correlation between model and (synthetic) empirical FC of *R*^2^ = 0.65. For reference, comparable modelling studies in the literature (including ones using brute-force fitting approaches), using similar data and parcellations, usually achieve *R*^2^ = 0.3 − 0.5 (Deco et al., 2013b; Abdelnour et al., 2014; Roberts et al., 2019). Panel C shows the distribution of errors (difference between estimate and target) in parameter estimates over all fit runs in the synthetic subject population. In these plots, error distributions that are centred on zero indicate a high level of accuracy of the fitted values, and error distributions that are narrow indicates a high level of consistency in parameter estimates across multiple synthetic subject runs, despite the algorithmic differences in terms of differing random initial conditions and differing patterns of random noise in the numerical integration routine. Recovery of the target values was excellent for three of the four neural dynamics parameters, with *g, g*^*EI*^ and *g*^*IE*^ showing errors of 1%, 10%, and 10%, respectively, and all three showing very high consistency over runs (narrow histograms). The *g*^*EE*^ parameter showed slightly poorer performance, with a broader range of estimated values over the synthetic subject population, but still with >75% of runs showing an error of <10% (lying within the range -0.1-0.1), and >99% showing an error of <20%. To demonstrate that the fit quality is not due to the large number of *C* parameters, this process is repeated in Figure S1, this time fixing the values in *C*, and only attempting to estimate *g, g*^*EI*^, *g*^*IE*^, and *g*^*EE*^. As can be seen, these parameters are recovered with similar accuracy in the reduced four-parameter model also.

**Fig. 3.**
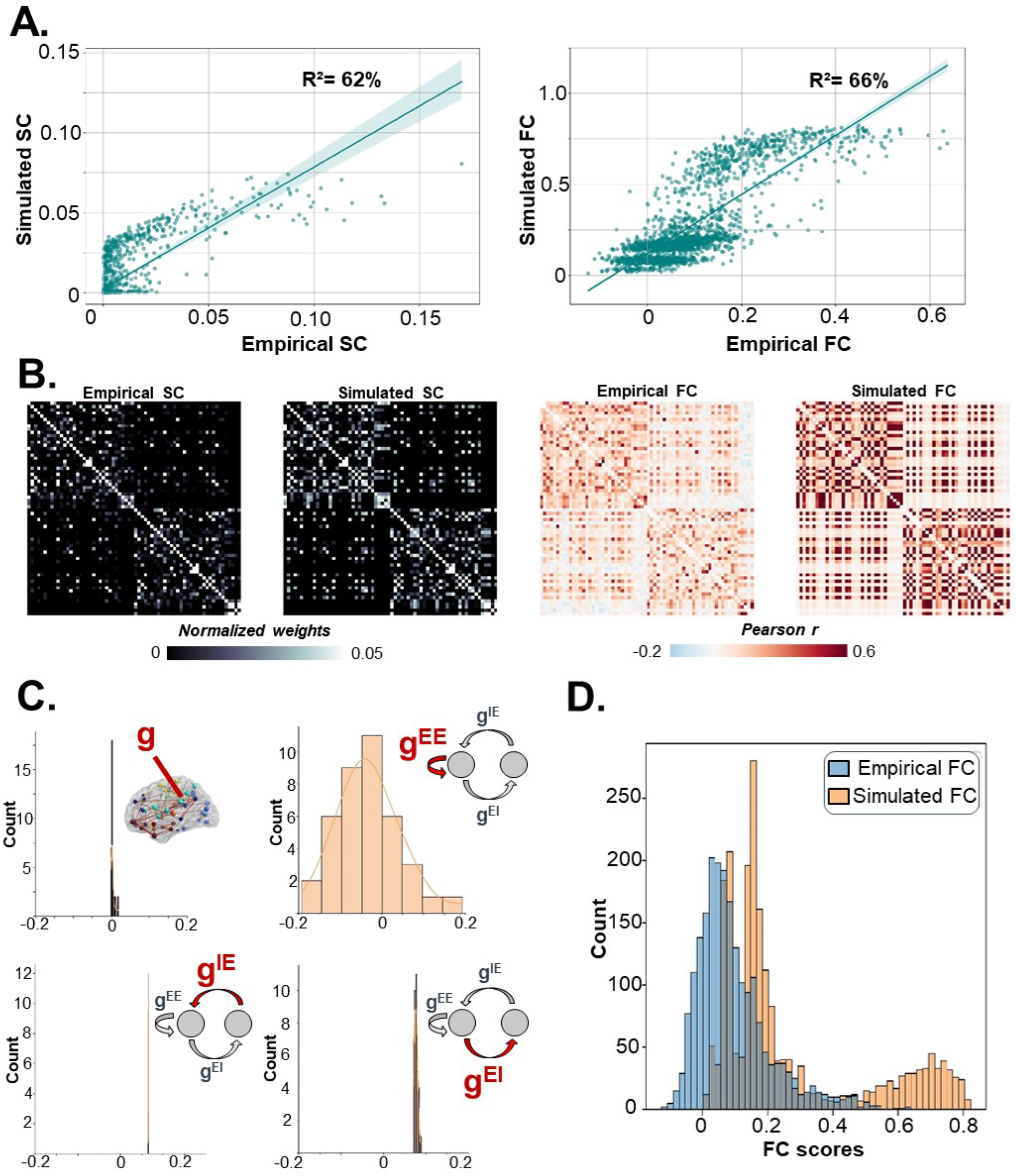
Synthetic Data Fitting Results. **A)** Scatter plots of (synthetic) ‘empirical’ anatomical connectivity (AC) and functional connectivity (FC) matrix values, against estimates of those quantities ‘simulated AC’, ‘simulated FC’, final fitted models, for a single exemplary subject. **B)** Same values from scatter plots in A), represented as AC and FC matrices. Note the accurately recovered left/right hemisphere block structure clearly present in both cases. **C)** Histograms of parameter estimate errors across the 50 synthetic subject fits for *g, g*^*EE*^, *g*^*IE*^, and *g*^*EI*^. In all 3/4 of these the width of the distribution is small, indicating reliable recovery of parameters across synthetic subjects. In one case (*g*^*EE*^), the range of errors is much higher than other three parameters, indicating potentially lower reliability in estimates of this parameter. **D)** Histogram of (synthetic) empirical and fitted FC values. Note the strong degree of overlap in the distributions, but with over-tendency towards slightly skewed and bimodal positive weight values in the simulated FC.

### 3.2 Fit results to empirical rsfMRI data

The model fitting performance for the 50 HCP subjects investigated in this study is shown in Figure 4. Similarly to Figure 3, Figure 4 Panel A shows empirical and model-estimated anatomical and functional connectivity matrices (in this case for three example subjects), and scatter plots of empirical vs. estimated FC. In all cases, the model-generated FC matrices showed excellent fits to the empirical FC data, with an average simulated-empirical FC correlation of *R*^2^=0.65, and a range of *R*^2^=0.4-0.9. This is a marked improvement on fits from comparable modelling studies in the literature (including ones using brute-force fitting approaches), using similar data and parcellations, which usually achieve *R*^2^=0.3-0.5 (Deco et al., 2013b; Abdelnour et al., 2014; Roberts et al., 2019). It is notable in the fitted FC matrices in Figure 4 that the secondary diagonals, representing strong FC between hemispheric homologues, are visible in the fitted FC matrices. This feature is robustly captured in FC measurements from fMRI and other sources, but is rarely captured in CNMM simulations to date (Deco et al., 2014), indicating our methodology’s ability to capture experimental data features that have been challenging for other techniques. As a further demonstration of the model’s ability to capture, and help understand known FC features, we used seed-based functional connectivity to identify default-mode network structures in empirical and simulated data. The way this is typically done with fMRI data is to use voxel-averaged time series from a region as a regressor in a Pearson correlation with voxel activity in the rest of the brain, with hot spots in the resultant map indicating high functional connectivity. Here, because we are using region-level data rather than voxel-level data, correlations across brain regions are shown as a pattern over ROIs on the cortical surface. Panel B of Figure 4 shows empirical (left) and simulated (right) correlation maps for seed ROIs in right precuneus, a canonical node of the so-called default-mode network (DMN), a constellation of correlated regions that is clearly evident in rsfMRI scans of most individuals. The surface maps in Figure 4B show cross-hemispheric correlations with the seed region as well as strong correlations with other DMN nodes in the lateral parietal and anteromedial frontal cortex.

**Fig. 4.**
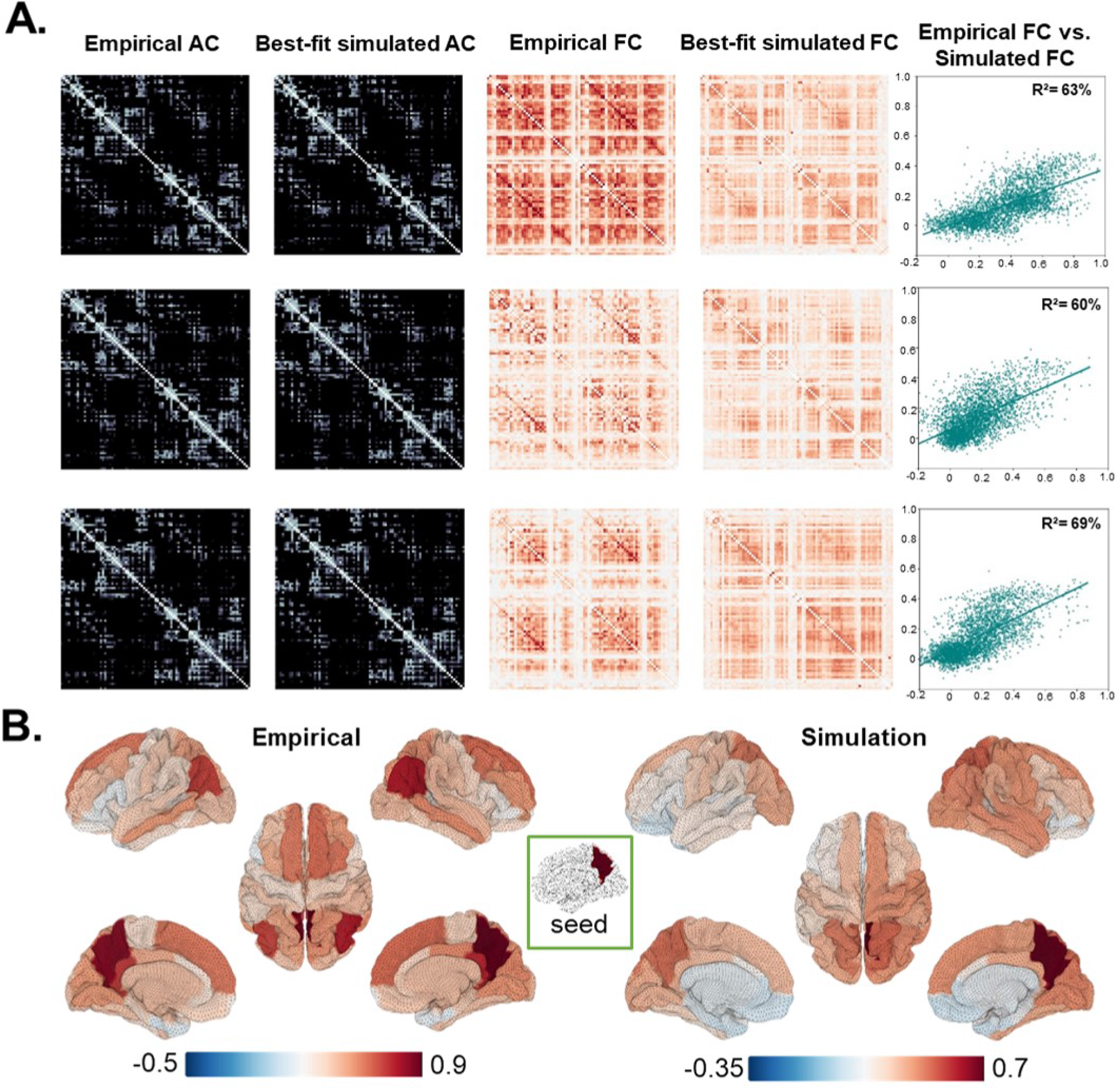
HCP Data Fitting Results - Example Subjects. **A)** For three example subjects (rows), from left to right: i) Empirical and estimated AC matrices, ii) empirical and simulated FC matrices, iii) scatter plots of empirical vs. simulated FC data from points. **B)** Empirical and simulated seed-based FC maps for a precuneus seed region for a single subject (from top row in panel A).

## 4 Discussion

In this paper we have outlined a novel methodology for estimation of neurophysiological model parameters using a deep learning-based computational architecture. Our focus has been on CNMMs, a particular type of neurophysiological model that typically focuses on approximately whole-brain connectivity and activity patterns, and are usually directed at understanding data from noninvasive neuroimaging modalities such as fMRI and M/EEG. As discussed later, the methodology has also proved highly effective for other CNMM applications such as EEG evoked potential waveforms (Momi et al., in prep) and neurocognitive models (Diaconescu et al., in prep), and the general approach should also lend itself easily to other model and data types in computational neuroscience and beyond.

Although the CNMM literature has expanded steadily over the decade or so since its inception, there has been relatively little technical work addressing the specific problem we have tackled here - namely parameter estimation for large brain network models. The problem is nevertheless an extremely important one, particularly if CNMM techniques are (as is surely desirable) to become usable and useful to the broader neuroimaging and neuroscience communities. Our approach builds on prior work of several groups, as well as introducing a number of novel components. In the following we discuss each of these, and in addition outline important limitations and potential future directions.

### 4.1 Methodological Highlights

Briefly, the methodologically novel aspects of our work are as follows:

#### Implementation

First, we have programmed the differential equations constituting the generative (neural dynamics) and measurement (haemodynamics) parts of a CNMM within one of the most widely used machine learning programming environments: PyTorch (Paszke et al., 2019). Whilst there are some syntactic and API-related differences between these implementations and the equivalent standard boilerplate Python code, they are fairly minimal. However, as we have emphasized, this implementation approach renders (albeit fairly implicitly) the CNMM as a computation graph, within which automatic differentiation can be used to calculate gradients of the objective function with respect to model parameters. In addition to the immediate functionality this provides in terms of optimization machinery, it is reasonable to expect that this general approach will increasingly be able to benefit from inevitable improvements in ANN-related software and hardware ecosystems in the coming years, much of which is centred around PyTorch and similar libraries.

#### Continuous-time approach and batching

Second, we use a time-continuous scheme where the model is run for 30 second increments, or *batches*. At the end of each batch, an update parameter value is chosen. A full run through the entire time series defines an *epoch*. At first glance this may seem problematic, since most neuroimaging scientists would likely not expect Pearson correlations computed over any given 30 second segments of a ten minute rsfMRI time series to closely resemble the Pearson correlations for the full time series. Nevertheless it does appear that the long-duration covariance structure is eventually learned by the model through the process of fitting multiple short-window batches successively. It is possible that the temporal variability in these short-window FCs actually facilitates the long-duration FC estimation, by providing multiple short excursions through a rich repertoire of brain states with slightly differing intrinsic covariance patterns. The issue that the windows are quite short is aided by the fact that parameter values from the previous batch are used to initialize the current batch, allowing the ADAM algorithm’s gradient descent to provide a ‘memory’ of sorts on parameters from earlier windows. This memory is reset however at the end of each epoch, at which point the final batch parameters are used to initialize the next sweep (of 30s batches) through the time series. This combination of *batching* and *epoching* - inspired by standard ANN methodologies - is a novel approach to CNMM parameter estimation that capitalizes on the fact that rsfMRI covariance structure has both strong trait-level properties that are across an entire session, as well as dynamics that vary slowly on the order of minutes Hansen et al. (2015). Sweeping through in batches thus allows a large number of iterations, allows the slow dynamics to inform the estimation process, as well as facilitating learning of the more stable and invariant aspects of the covariance structure.

#### Bifurcation analyses

Third, we undertook a bifurcation analysis to find constraints on the RWW model parameter space, allowing us to restrict the system to a regime showing more neurobiologically realistic dynamics, and that also yields superior fit results than other sections of the parameter space. On the basis of this, we were able to restrict the sampling of the *gEI* and *gIE* parameters to combinations lying on a polynomial spline function, thereby collapsing a 2-dimensional parameter space onto a 1-dimensional manifold within that space (Figure 2B).

### 4.2 Key Results

#### Accurate recovery of CNMM model parameters

We demonstrate the performance of our approach by applying it to synthetic (Section 3.1) and empirical (Section 3.2) data. The synthetic rsfMRI data results demonstrate accurate recovery of model parameters in a situation where the ground truth is known. The anatomical connectivity matrix used to generate synthetic data was the widely-used 66×66 matrix of (Hagmann et al. 2008; see also e.g. Deco et al. 2013b). Here, we showed that the four neural dynamics parameters can be recovered with near-zero errors. We then applied the approach to rsfMRI data from the Human Connectome Project. Our model showed a good ability to recover the relatively strong FC between some pairs of homologous regions in opposite cerebral hemispheres. This pattern which manifests in the connectivity matrices as a secondary diagonal in the upper right and lower left quadrants - is an extremely robust feature of FC measurements. Moreover, it is also a pattern that has historically been rather poorly reproduced in CNMM simulations (Deco et al., 2013b, 2014). As shown in Figure 4 panel B, a closer look at individual columns of the simulated FC matrices show good qualitative correspondence with empirical FC data, when viewed as ROI weightings over a brain surface. For example, familiar FC patterns such as the default mode network structures that are strongly correlated with the precuneus seed can be clearly seen in these visualizations.

#### Number of parameters

One of the important advantages of model optimization approaches, as compared to the often used brute-force approach discussed earlier, is the ability to fit more than a handful of parameters. We have demonstrated this here by exploring two sets of models in our analyses of synthetic data, one with a small number of estimated parameters, and one with a very large number. The first of these focused on just the the four RWW parameters *g*^*EE*^, *g*^*EI*^, *g*^*IE*^, and *g*, and yielded excellent fit results (Figure S2). The second set of synthetic data analyses these same four parameters, and in addition fitted the roughly 1000 elements of the anatomical connectivity weights matrix *C* (Figure 3). Importantly, as can be seen by comparing Figures 3 and S2, both of these yielded excellent results, with similar error levels for the RWW gain parameters in both cases. Estimation of (effective) connectivity weights (entries of *C*), as is done here, has been the principal functionality of small-scale neural mass modelling approaches, such as with DCM. In contrast, CNMM studies have usually opted to keep anatomical connectivity fixed (by the structural connectome), and focus instead on estimating global neural dynamics parameters, of the kind studied here. Connectivity weight estimation has been performed for large networks in some recent work (Deco et al., 2013b; Shen et al., 2019; Singh et al., 2020; Frässle et al., 2020), but this remains an under-developed area of study.

#### Noise-driven brain dynamics

As noted above, when the entire CNMM is specified programatically as a computation graph, it resembles very closely a deep neural net, that can be understood as propagating ‘patterns’ from the input layer, through the hidden states, on to the output layer. Because a variety of computational steps can be encoded throughout the graph, we can specify either the simulated time series, or the simulated FC, as the output layer. For sensory- or stimulation-evoked neural dynamics, the sensory or electromagnetic stimulus would be used as the input and defined as an input layer. For resting state neural dynamics there is still an input layer, but the input is random noise injected into all nodes through the noise term *σv*^*E/I*^ in Eqs. 4-5. This noise signal is propagated through the computational graph, and is converted into structured temporal fluctuations and covariance patterns on the output layer. This is in keeping with one of the general current perspectives in CNMM work, namely that resting state functional connectivity patterns reflect random fluctuations (uncorrelated background spiking activity) that are filtered by endogenous brain dynamics and connectivity structure (Deco et al., 2013b, 2014).

### 4.3 Assumptions and limitations

Our framework has a number of assumptions and limitations that should be examined closely in future research looking to build on the present work.

#### Other models

Future work should abstract and generalize the model-fitting methodology used here in such a way that it could be easily deployed for different models of neural dynamics. Indeed, the two aspects of our implementation (the analytic Laplacian and decision to constrain the parameter space of the RWW model based on the bifurcation analysis considerations) make the results shown here highly specific to both the model and dynamic regime of interest. Future work building on this specific model could consider whether these assumptions on the model type etc. can be relaxed, for comparable performance.

### 4.4 Relation to previous work

Although, as we have indicated, the prior literature on parameter estimation for CNMMs is relatively small, both this problem and related ones (such as parameter estimation for small networks) have been studied previously by several groups. Most notably, the approach of specifying CNMMs within an optimization framework that employs gradients computed via AD was (to our knowledge) introduced by the group of Jirsa and colleagues, in the context of their ‘Virtual Epileptic Patient’ (VEP) work (Proix et al., 2017; Jirsa et al., 2017; Hashemi et al., 2020). These authors used the STAN library, which is an older and more established tool than PyTorch, and that has for some time been leading the development of ADVI (AD-based Variational Inversion) techniques, with applications across a variety of scientific fields. The principal focus of this VEP work is to use a CNMM developed specifically to model epileptic seizure spreading (Jirsa et al., 2017) to identify alternative and potentially superior targets for epilepsy - surgery that may not have been considered with conventional assessment methods. Specifically, ‘epileptogenicity’ parameters are estimated for all nodes in a medium-sized or large-scale anatomical brain network, based on seizure recordings from implanted stereotactic EEG (sEEG) electrodes. In one of the most recent iterations of this work, Hashemi et al. (2020) have developed MCMC sampling methods that pair with ADVI-derived gradients and numerical integration to give an efficient and robust probabilistic inference approach to Bayesian inversions of Epileptor models from individual patient data.

The use of the Variational Bayesian techniques for physiological model parameter estimation was first introduced in 2003 by Friston and colleagues under the name ‘Dynamic Causal Modelling’ (DCM; Friston et al. 2003), developed extensively by that group over the subsequent 20 years. The ‘classic’ DCM treats the brain as a deterministic nonlinear dynamical system, with parameters mainly describing inter-regional connectivity weights within a small network of brain regions. The connectivity parameters are estimated by ‘perturbing’ the system (typically with a sensor or cognitive task) and measuring the transient neural activity response that follows. To measure the first-order connectivity between brain regions in the absence of input, the matrix of connectivity weight parameters (which for a first-order linear system, is the system’s Jacobian) is estimated from a set of input time series. The ‘standard’ DCM model fitting scheme is based on gradient descent on the log model evidence, using analytically-defined or numerically estimated (not AD-derived) gradients. When performing the estimation for parameters and hyperparameters during the expectation-maximization step, the Fisher scoring ascent, a type of gradient ascent, of the objective function uses the log posterior of the likelihood and prior potentials (Friston et al., 2003). The log posterior of the DCM is computed using likelihood and prior terms, with the former based on Gaussian assumptions about the errors in the observation model of the DCM. More recently, sampling-based approaches for DCMs have also been proposed and implemented (Sengupta et al., 2016). For the most part, this approach has been limited to parameter estimation for small network models (Gilson et al., 2016); although more recent work has extended DCM for fMRI to larger networks of dozens to hundreds of nodes (Frässle et al., 2020). In both these cases (small and larger fMRI DCMs), a notable difference as compared to the present work, and CNMMs in general, is the time scale of the generative model. In our model, as with earlier work (e.g. Deco et al. 2013b; Demirtaş et al. 2019), the model of neural dynamics evolves on the timescale of milliseconds, commensurate with known time scales of single neuron and neural population activity. Many neural mass models for fMRI (Cabral et al., 2011; Gilson et al., 2018; Friston et al., 2003; Abdelnour et al., 2014), including fMRI DCMs, instead have dynamics on the order of seconds, which is actually commensurate with timescale of BOLD fMRI data. Slow timescale models are, as a rule, easier to work with as they require less run time and have less complex generative equations, with the disadvantage that they are less physiologically accurate.

### 4.5 Conclusions and Future Directions

The model and methodology proposed in this paper represent our first attempt at applying the relatively new, but rapidly developing, parameter estimation functionality of modern machine learning software libraries to our use case of parameter estimation for neurophysiological models. Although we have found excellent performance in the test cases examined, we still consider this work to be largely a proof-of-principle and a demonstration of feasibility. Future work can build on the methodology presented here in a number of ways.

#### Model types

There is considerable scope for generalization of our implementation to accommodate a wider range of model types and specifications. Currently in the code base accompanying this paper (github.com/dl-paramest-for-neurophys-models), the WWD model is hard-coded. This can however be easily generalized. Other systems of equations that have been used in a CNMM context for fMRI include Kuramoto (Cabral et al., 2011), Stefanescu-Jirsa (Falcon et al., 2015), and Linear Diffusion (Abdelnour et al., 2014; Robinson, 2012) models. Several elements of the approach presented here (specifically the short-window batching approach and utilization of AD and ADAM) should be broadly applicable to most if not all model types. Other elements such as the parameter constraints based on bifurcation analyses of the RWW equations are less directly transferable, although an analogous approach could easily be applied in many cases with some additional assumptions and derivations. Similarly, our approach could be generalized to other data types and experimental contexts, including task fMRI, sensor- and source-space MEG/EEG/fNIRS, and noninvasive brain stimulation. In particular, the overall methodology works as well if not better than for rsfMRI when applied to fast-timescale evoked responses to transcranial magnetic stimulation (TMS) pulses as measured by concurrent TMS-EEG (Momi et al., in prep). Building on this and on the above demonstrations, another focus of our current work is the fitting of multimodal models, which jointly describe diverse concurrently recorded signals (e.g. concurrent EEG-fMRI or concurrent EEG-fNIRS).

The ability of our methodology to capture FC patterns previously poorly reproduced in CNMM simulations implies that the resulting models might be more appropriately applied to the wide range of computational questions that require whole-brain simulations. As just one example, a variety of computational techniques have been applied to the study of seizure propagation throughout the brain (Olmi et al., 2019; Liou et al., 2020; Kramer et al., 2005), a topic of vital importance to improving our understanding of epilepsy. CNMMs generated using our methodology are well suited for continued study of seizure propagation using whole-brain models constrained directly from human data, and can potentially be synthesized with the growing number of biophysically-detailed human neuron models (Rich et al., 2021; Beaulieu-Laroche et al., 2018; Kalmbach et al., 2018) towards a better understanding of the multi-scale nature of seizure propagation.

#### Simulator libraries

Another dimension to the question of generalization and generalizability is at the implementation and software level, and pertaints to the distinctions outlined earlier between black-box and glass-box parameter estimation approaches. There are a number of existing and well-developed CNMM simulator libraries currently available (e.g. TVB (Ritter et al., 2013; Sanz-Leon et al., 2015), SPM-DCM, Brain Dynamics Toolbox (Heitmann et al., 2018), PyRates Gast et al. (2019), NeuroLib Cakan et al. (2021)), that allow simulations not only with the RWW equations but a wide variety of alternative models for neural dynamics. Unfortunately however, the ‘glass-box’ methodology described here would be either difficult or impossible to directly deploy in most if not all of these cases, because the code implementations to not support AD-computed gradients. This is for a variety of reasons, including separated front-end and back-end code execution / generation (PyRates), optimized C-code (TVB) or Numba jit-compiled (NeuroLib) backends. An interesting broader question here is whether greater integration with explicitly ML-oriented programming environments are a natural fit neurophysiological modelling applications, or alternatively whether the relevant computational and algorithmic infrastructure will simply become part of the ‘boilerplate’ general scientific computing stack.

#### Conclusion

It is highly likely that the coming years will see a greater integration of machine learning tools in computational neuroscience applications, including physiological modelling of neuroimaging data using CNMMs. We expect that this will greatly facilitate progress in estimation of CNMM parameters, and deployment of CNMMs in clinical and other practical contexts. The approach outlined in the present paper points to some promising new directions forward for improved computational tool development in this area.

## 5 Acknowledgments

JG acknowledges research funding support from the Krembil Foundation, CAMH Discovery Fund, and Tri-Council UK-Canada AI Initiative.

## Footnotes

1 Note that the *objective function* Jacobian, defining a matrix of partial derivatives of model parameters with respect to the objective function, is different to Jacobians that might define or aid a numerical integration routine used for simulating neural activity, which would be a matrix of partial derivatives of the system *states* with respect to each other. This latter *neural dynamics Jacobian* could nevertheless be, and often is, used in manual calculation of an analytic optimization Jacobian by application of the chain rule.

