## Supplementary Materials for "Deep Learning-Based Parameter Estimation for Neurophysiological Models of Neuroimaging Data"

PREPRINT

May 19, 2022

### A. Model Fitting and Batching Schematic

An important part of our model fitting approach is to break up the rsfMRI time series into short windows that serve as *batches* in the machine learning parlance. This process, described in the main text, is summarized graphically in Figure S1 below.

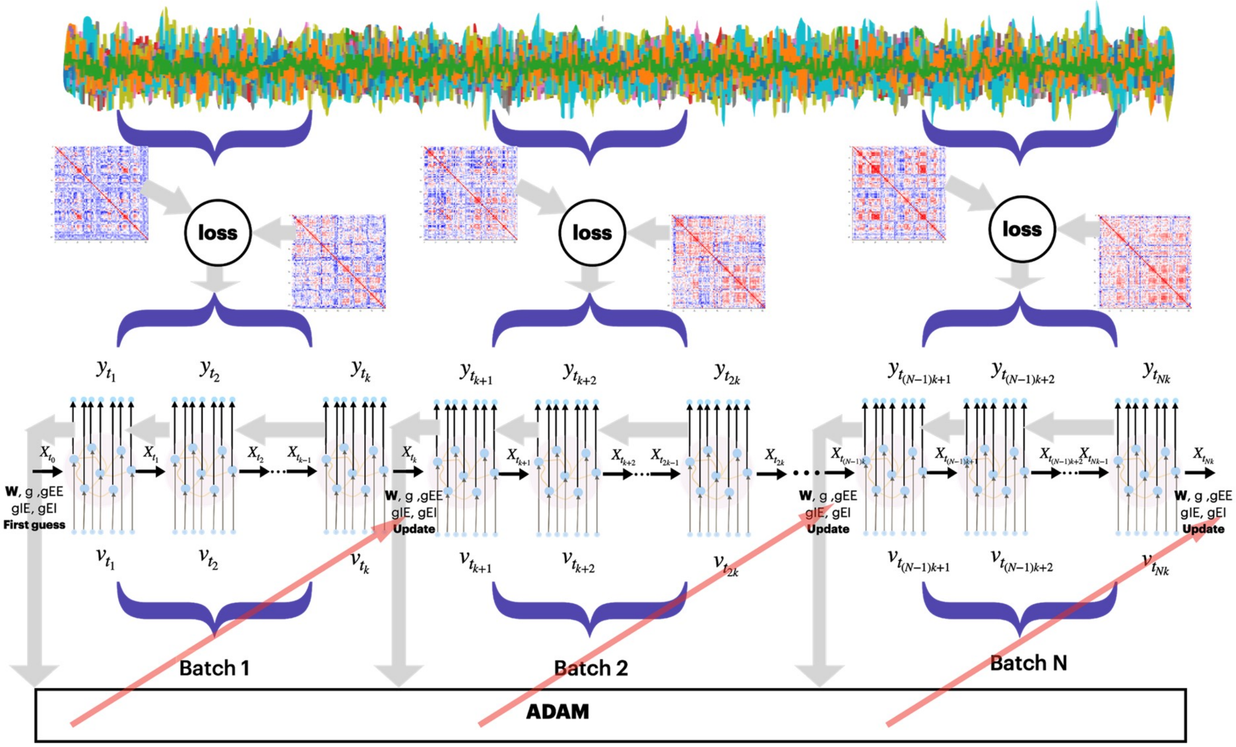


**Fig. S1 | *Model and model fitting schematic.*** The model fitting process is divided into *batches* (30s segments of data over which the model is run, loss is computed, and parameters are updated), and *epochs* - full runs through the entire time series, to be repeated until convergence. For each batch, the objective function uses the input data to minimize the free energy (a complexity-penalized model fit term) of the simulated states given the data and model parameters. Physiologically realistic neural activity and BOLD functional connectivity patterns are maximized near the critical (bifurcation) point of the bistable regime. The backpropagation step then obtains the gradient (loss) which is used to update the model parameters ($C$, $g$, $g^{EE}$, $g^{EI}$, $g^{IE}$) such that the states of the RWW model can be updated, and a batch of BOLD signals can be produced. A haemodynamic forward model returns BOLD signals at each element in the batch, from which the Pearson correlation FC is calculated and used in the objective function. The PyTorch ADAM optimizer is then used for optimizing the parameters.

### B. Model parameters

A full list of the parameters neural, haemodynamic, and optimization parameters, including nominal values, are given in Table S2.

| SYMBOL | NAME | UNIT | NOMINAL VALUE |
|---|---|---|---|
| **Neural Model** | | | |
| $E_i$ / $I_i$ | Average synaptic gating variable at local area $i$ | n/a | n/a |
| $R^E$ / $R^I$ | Population firing rate, converted from $E/I$ | n/a | n/a |
| $I^E$ / $I^I$ | Input currents to $E$/$I$ neural populations | n/a | n/a |
| $a$ | Parameters of the input-output function | n/a | (E) 310, (I) 615 |
| $b$ | Parameters of the input-output function | kHz | (E) 125, (I) 177 |
| $d$ | Parameters of the input-output function | ms | (E) 0.16, (I) 0.087 |
| $w$ | Local excitatory recurrence | n/a | 1.0 |
| $\gamma$ | Kinetic parameter | ms | (E) 0.641, (I) 1 |
| $\tau$ | Kinetic parameter | ms | 100 |
| $J_n$ | Synaptic couplings | nA | 0.2609 |
| $I_0$ | Overall effective external input | n/a | n/a |
| $C_{ij}$ | Entries of the anatomical connectivity matrix | n/a | 80. |
| $L$ | Anatomical SC Laplacian | n/a | n/a |
| $G$ | Global scaling coupling (reweighs the SC) | nA | n/a |
| $\eta$ | Amplitude of Gaussian white noise | nA | 0.01 |
| $\sigma$ | Standard deviation of Gaussian white noise | n/a | 0.02 |
| $g^{EI}$ | Within-node excitatory-to-inhibitory synaptic gain | n/a | 0.501 |
| $g^{IE}$ | Within-node inhibitory-to-excitatory synaptic gain | n/a | 0.4774 |
| $g^{EE}$ | Within-node excitatory-to-excitatory synaptic gain | n/a | 2.49 |
| $g$ | Long-range global coupling | n/a | 82 |
| $\lambda$ | Scaling of long-range feedforward inhibition | n/a | 0 or 1 |
| $I_0$ | Constant external input | n/a | Varied |
| **Haemodynamic Model** | | | |
| $F$ | Volume flow rate into the tissue | ml/s | n/a |
| $Q$ | Normalized total deoxyhemoglobin content | n/a | n/a |
| $V$ | Normalized venous volume | n/a | n/a |
| $X$ | Flow between different steps of the model | n/a | n/a |
| $Y$ | BOLD signal | n/a | n/a |
| $\rho$ | Haemodynamic rate constant | n/a | 0.34 |
| $\tau 0$ | Haemodynamic transit time | n/a | 0.98 |
| $\alpha$ | Haemodynamic stiffness | n/a | 0.32 |
| $\tau s$ | Haemodynamic signal decay | n/a | 0.65 |
| $\tau f$ | Haemodynamic autoregulation | n/a | 0.41 |
| $k$ | Constant coefficient | n/a | n/a |
| **Parameter Optimization** | | | |
| $\Theta$ | All model parameters | n/a | n/a |
| $O$ | Objective Function | n/a | n/a |
| $c_{a/b/c}$ | Optimization hyperparameters | n/a | n/a |

**Table S1 |** ***Neural, haemodynamic, and optimization model parameters and variables.***

#### C. Additional parameter estimation results

Further results are shown that continue the synthetic data analyses in Figure 3. Figure S2 shows model fit results for four-parameter version of the model, which is identical to the model described in the main text but with the connectome matrix weights $C$ are removed. As can be seen, the parameter estimate error distributions over 50 synthetic subject runs return the same results as do the full model. Panel B of Figure S2 additionally shows posterior correlations of model parameters. As should be expected, $g^{IE}$ is strongly anti-correlated with $g^{EI}$, these being antagonistic forces within the model equations. Panel C demonstrates model converge over runs. The degree of convergence of a random Markov Chain can be estimated using the Gelman-Rubin convergence statistic $\hat{R}^2$, based on the stability of outcomes between and within chains of the same length. Values close to one indicate convergence to the underlying distribution, and values greater than 1.1 indicate inadequate convergence. The $\hat{R}^2$ values computed for the synthetic data model runs, shown in Figure S2 Panel C, indicate good convergence performance of the model.

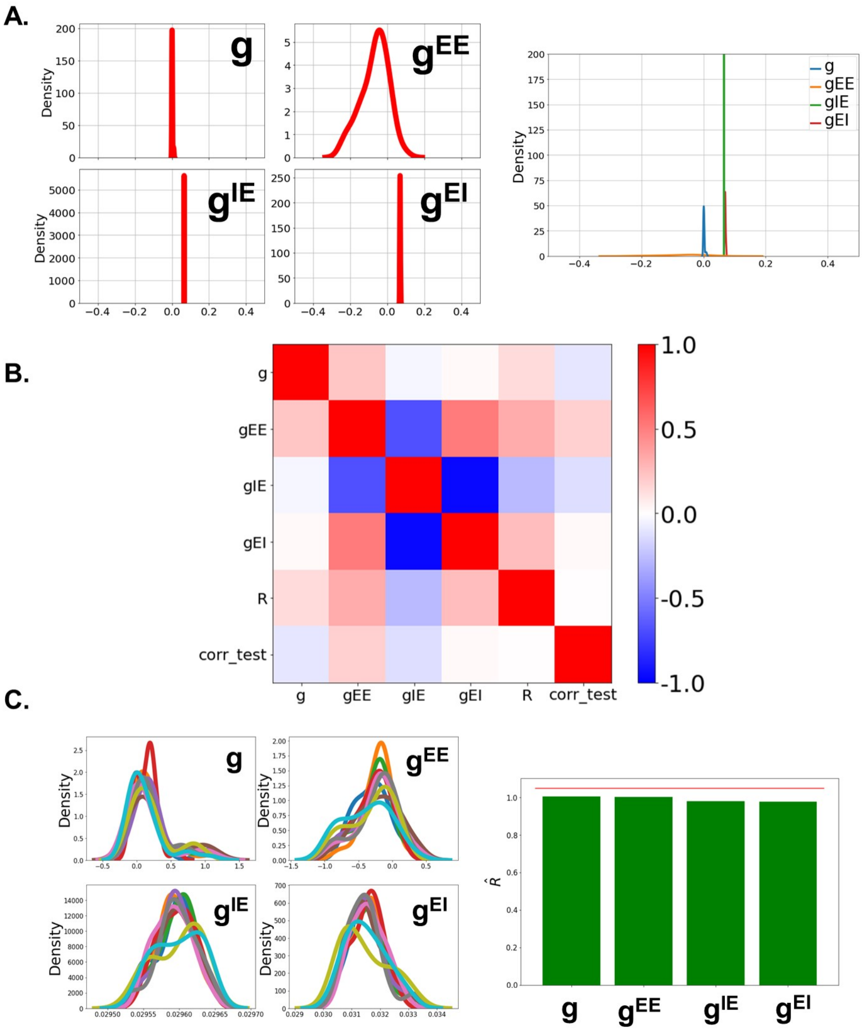


**Fig. S2 | *Four-parameter synthetic model fitting results.*** **A**. Error distributions for the four RWW parameters over the 50 synthetic subject model runs. Note that these are virtually identitical to the corresponding distributions in Figure 3, despite the additional $C$ parameters estimated in that case. **B**. Posterior correlation matrix of model parameters. **C**. Posterior parameter distributions ove rmodel runs (left) and Gelman-Rubin convergence statistic $\hat{R}^2$ (right)
